# Effect of Stroma-directed Drugs in Combination with Chemotherapy Against Pancreatic Cancer- a Preclinical Study

**DOI:** 10.1101/2024.12.24.630138

**Authors:** Mamta Gupta, Hoon Choi, Emma E Furth, Sydney Shaffer, Stephen Pickup, Cynthia Clendenin, Fang Liu, Quy Cao, Hee Kwon Song, Yong Fan, Jeffrey Duda, James C Gee, Thomos Karasic, Mark Rosen, Peter O’Dwyer, Rong Zhou

## Abstract

Cytotoxic chemotherapy plays an important role for extending the survival of patients with pancreatic ductal adenocarcinoma (PDAC). To enhance the efficacy of chemotherapy for eradicating the cancer cells, we have compared the standard care chemotherapy (combination of nab-paclitaxel, gemcitabine and cisplatin, NGC) versus NGC plus stroma-directed agents (calcipotriol and losartan, respectively) in a genetically engineered mouse model of PDAC. Over a 2-week study period, MRI was conducted to measure the tumor size and to test the sensitivity of imaging markers derived from diffusion-weighted imaging (DWI), dynamic contrast enhanced MRI (DCE) and magnetization transfer ratio (MTR) for assessing the tumor cellularity and stromal changes. Detailed immunohistochemistry and preliminary single cell RNA sequencing (scRNAseq) study were applied to tumor tissues collected upon euthanasia on day-14. Our major findings are: 1. Compared the untreated controls, NGC chemotherapy induced significant tumor growth inhibition and stromal changes including pronounced reduction of fibroblast associated protein (FAP) level accompanied by increased matrix collagen content, significantly reduced microvascular permeability revealed by DCE corroborated with reduced microvascular density. 2. Losartan+NGC significantly enhanced inhibition of tumor growth beyond NGC and increased lymphocytes infiltration in the tumor which may contribute to enhanced cancer cells eradication. 3. NGC treatment enriched the fraction of mesenchymal (M) subtype while reducing the epithelial (E) subtype of cancer cells compared to the controls, and this trend was reversed by calcipotriol+NGC. In conclusion, our study captured changes in cancer cell and tumor microenvironment in response to chemo stromal therapy versus chemotherapy alone with mechanistic insights.

## Introduction

Pancreatic ductal adenocarcinoma (PDAC) is an aggressive and lethal form of cancer with limited treatment options. Systemic chemotherapy remains an important treatment for almost all PDAC patients as neoadjuvant or adjuvant therapy for non-metastatic disease and as first line therapy for metastatic disease (1–4). The successful implementation of multidrug chemotherapy regimens such as gemcitabine /nab-paclitaxel /cisplatin and FOLFIRINOX (leucovorin calcium, fluorouracil, irinotecan hydrochloride, and oxaliplatin) has been the driving force that prolongs the survival of PDAC patients and the standard care treatment of PDAC. Chemotherapy alone however cannot eradicate the cancer due to intrinsic and extrinsic factors. Intrinsically, sensitivity to chemotherapy is influenced by the subtype of cancer cells since epithelial (or classical) subtype is relatively sensitive to chemotherapy whereas mesenchymal or quasimesenchymal subtype more resistant (5–9). Extrinsically, the dense stroma as a unique histological feature of PDAC, harbors a tumor microenvironment where extracellular matrix proteins and stromal cells cooperate to impede drug delivery, reduce the cytotoxicity of chemotherapy drugs, and sustain cancer cell survival (10–12). Stroma-directed agents, for example, synthetic vitamin D such as calcipotriol /paricalcitol has been shown to reprogram the stroma phenotype, leading to increased efficacy of gemcitabine in murine PDA model (13).

Losartan, a clinical medication for management of high blood pressure has been shown to increase the efficacy of FOLFIRINOX plus chemoradiation treatment of locally advanced PDAC patients, leading to downstaging of their tumors for surgical resection (14,15). Hence combination of chemotherapy with stroma-directed agents may represent a safer approach to enhance the efficacy of chemotherapy without significantly increasing the toxicity of the treatment. The long time course (several months) of chemo or chemo stromal therapy would benefit from minimally invasive markers such as those obtained from blood plasma or *in vivo* imaging to help physicians evaluate the treatment and make decisions, e.g., escalating, deescalating or switching to a different treatment if the current one is ineffective.

Since CT and MRI are standard imaging modalities for diagnosis and staging of PDAC in the clinic and MRI offers superior soft tissue contrast, we have conducted an MRI-oriented preclinical study using a genetically engineered mouse model (GEMM) of PDAC. According to study design in **Figure 1**, we compared the standard care chemotherapy consisting of nab-paclitaxel, gemcitabine and cisplatin (NGC) with NGC+X (X = calcipotriol, losartan) and untreated controls. We hypothesized that MRI-based tumor size provides an objective assessment of the treatment effect per RECIST criteria (16), meanwhile other clinically applicable MRI methods, including dynamic contrast enhanced MRI (DCE), magnetization transfer ratio (MTR) and diffusion-weighted MRI (DWI) can detect treatment responses in stroma, which has three major components, namely microvasculature, extracellular matrix (ECM) and cancer-associated fibroblasts (CAFs) (17). In the past, we and others have shown the utility of DCE to detect vascular effect mediated by stroma directed drug (18) and chemotherapy (19) in PDAC, DWI to evaluate PDAC tumor cellularity (20) and change in response to treatment (21). MTR appears to be sensitive to increased matrix collagen deposition in liver fibrosis and PDAC (22,23), however, MTR signal could be dominated by cell density change (24). Our study design includes detailed immunohistochemistry (IHC) analyses to assess key stromal components and for verifying the MRI metrics, for example, CD31 to estimate MVD and Sirius red to estimate matrix collagen level and fibroblast activation protein-α (FAP) for treatment response since FAP, as a CAF signature, is associated with the survival of PDAC patients (25) . Furthermore, preliminary single cell RNA sequencing analyses were conducted to evaluate potential alterations of the molecular subtype of cancer cells and to provide mechanistic insights based transcriptome changes for mechanistic insights and generation of hypothesis for further studies.

**Figure 1.**
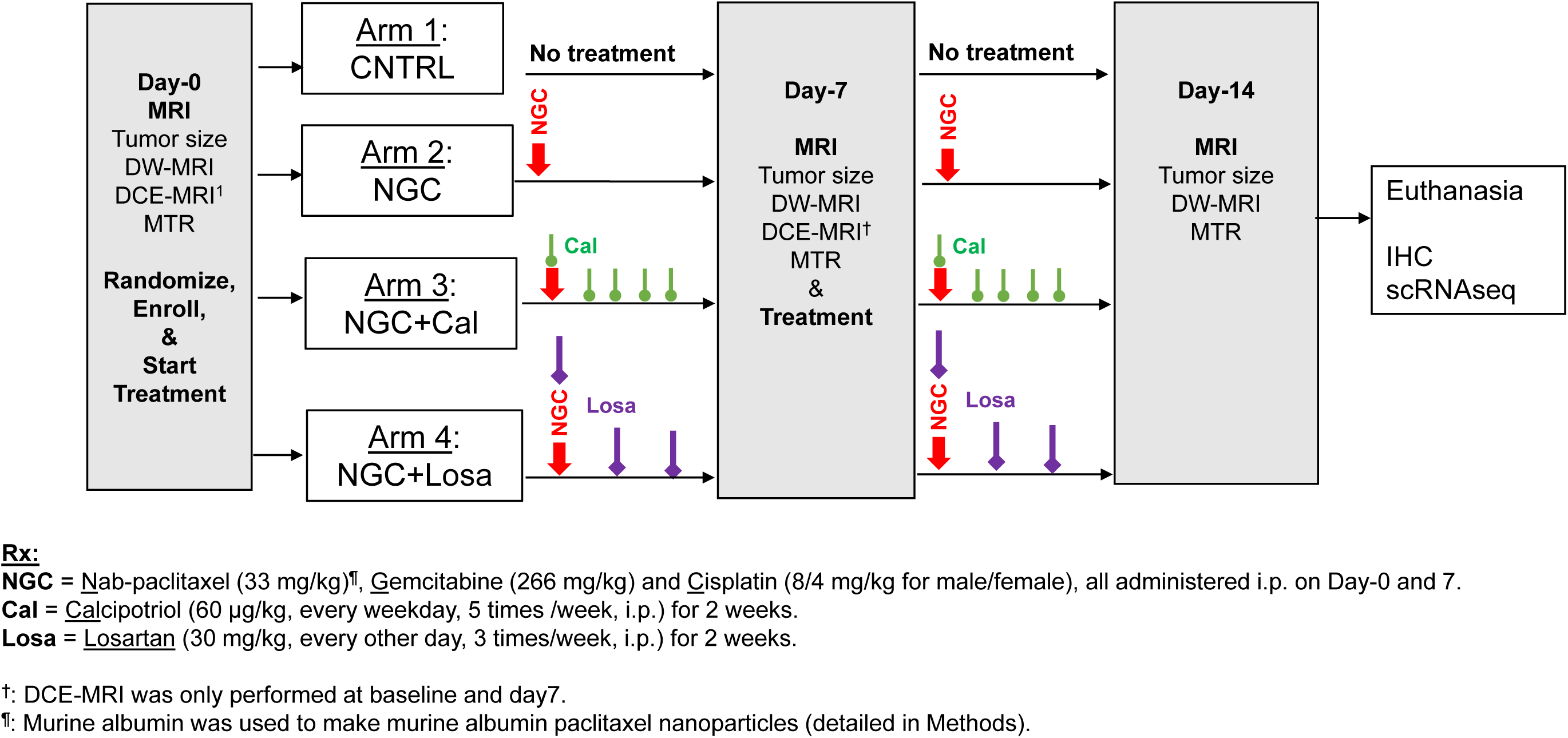
Design of pre-clinical trial. The preclinical trial lasted for 2 weeks, and all surviving mice were euthanized on Day-14. Tumor size was measured in all mice enrolled on Day0, 7 and 14 by T2W MRI. DW-MRI and DCE-MRI protocols were applied to a subset of mice due to limited scanner availability and DCE-MRI was further limited by the requirement of tail vein catheterization hence only mice having two successful catheterizations (on Day0 and 7) contributed data.

## Materials and Methods

### Materials

Gemcitabine, cisplatin, losartan and ProHance®, a Gd-based contrast agent used in the clinic were all purchased from the Pharmacy of the Hospital of University of Pennsylvania (HUP).

Calcipotriol, which has been administered to mice (13), is a synthetic derivative of calcitriol (1,25 OH-cholecalciferol), a biologically active form of vitamin D. Pharmaceutical grade of calcipotriol (catalog 270010) and murine serum albumin (catalog 502036442) were purchased from Fisher Scientific (Waltham, MA, USA), and paclitaxel (Catalog P-9600) from LC Laboratories (Woburn, MA, USA).

### Manufacturing murine version of nab-paclitaxel using murine albumin

The protocol which was used for preparation of nab-paclitaxel (26) was modified by replacing human albumin with murine albumin to make the murine version of nab-paclitaxel. Briefly, 30 mg of paclitaxel was dissolved in 0.55 ml of chloroform and 0.05 ml of ethanol. The solution was then added to 30 ml PBS solution containing 1% w/v murine albumin, which was pre-saturated with 1% chloroform. The mixture was transferred into a high-pressure homogenizer (EmulsiFlex-C5, Avestin).

Emulsification took place at pressures around 15,000 psi with the emulsion being recycled for a minimum of 6 cycles. The solution was then moved into a rotary evaporator to remove the chloroform and subsequently lyophilized (Labconco™ FreeZone™ 4.5L, Fisher Scientific). The lyophilized powder was stored at -20°C until use. The powder was reconstituted in PBS to 5 mg paclitaxel /ml and was injected in mice shortly after preparation.

### In vivo studies

All animal procedures were approved by the institutional animal care and usage committee (IACUC) of the University of Pennsylvania.

KPC mice (Kras^G12D^:Trp53^R172H^:Pdx1-Cre) were bred at the Mouse Hospital of the Pancreatic Cancer Research Center, the University of Pennsylvania. Screening for tumors was done via weekly abdominal palpation starting at 11 weeks of age followed by ultrasound examination (Vevo 2100, VisualSonics, Toronto, ON, Canada).

**Study design** is shown in **Figure 1**. Both male and female mice were transferred from the Mouse Hospital and were randomized to enroll in the treatment groups. Study design features: 1) a chemotherapy backbone consisting of murine version of nab-paclitaxel (detailed in Materials), gemcitabine and cisplatin (NGC); 2) stroma-direct agents including synthetic vitamin D (calcipotriol, Cal) and losartan (Losa), 3) a 2-week trial where NGC chemotherapy was administered on Day-0 and 7 with stromal drugs given more frequently indicted in Figure 1. T2W, DWI and MTR were conducted at Day-0 (before treatment), 7 and 14 while DCE-MRI performed on Day-0 and 7.

#### MRI acquisition

MRI studies were performed on a 9.4 T Avance III console (Bruker, Berillica, MA, USA), equipped with 12 cm ID, 40 G/cm gradients. To prepare animals for MRI exams, general anesthesia was induced and maintained via free breathing of 1–3% isoflurane in oxygen through a nose cone. A rectal temperature probe and pneumatic respiration pillow (SAII, Stonybrook, NY, USA) were applied to the animals and the animals were positioned on a 3D printed bed, which was positioned in a 35 × 40 (ID × length) mm quadrature birdcage coil (M2M, Cleveland, OH, USA). The RF coil was positioned in the magnet and regulated warm air source was directed over the animals to maintain core body temperature at 37±0.2°C. Respiration rate and core body temperature were monitored throughout the MRI exam.

Following calibrations of the scanner and generation of scout images, an axially oriented contiguous series of T2W images spanning the entire tumor was generated using a TurboRare protocol as described earlier (21). The image orientation and abdominal region coverage of the T2W scan were passed to DWI, DCE and MTR scans.

To mitigate respiratory motion-induced artifacts in DWI, a radial k-space sampling scheme (19,27) was employed with acquisition parameters of diffusion time (Δ) = 14 ms, diffusion gradient duration (δ) = 9 ms, b-value = 10, 535, 1070, 1479, and 2141 s/mm^2^, TE/TR = 28.6/750 ms, matrix size = 64 × 403, averages = 1, bandwidth = 50 kHz with fat saturation and total acquisition time approx. 25 min. No respiration gating was employed while the mouse was breathing freely during acquisition.

For DCE MRI a tail vein catheter was placed with an extension tubing of sufficient length to reach from the magnet isocenter to the end of the magnet bore. The tubing was preloaded with contrast agent in order to minimize dead volume effects. To enable optimal special and temporal resolution of DCE, 3D stack-of-stars (SoS) sequence was applied to all acquisitions in the DCE protocol including B1 maps by actual flip angle imaging (AFI), T1 maps by variable flip angle (VFA) imaging and the DCE series by a golden-angle (111.25°) ordering scheme to obtain a uniform coverage of k-space we described previously (28). Two minutes after starting the DCE series, 0.2 mL of diluted Prohance was injected over 10 s manually via the catheter. All components of the DCE protocol had matrix size = 64×201×16 and FOV = 32×32×8 mm^3^ as we described earlier (21).

For MTR, a magnetization prepared 3D GRE was acquired with FOV= 32x32x8 mm, matrix= 64x64x16, TR/TE /flip = 5.7 /2.8 ms /5°, averages = 4 with an 18 sec 2.5 µT saturation pulse applied at 4 kHz (*MT_on_* image) and 250 kHz off-resonance (*Reference* image), respectively. Each saturation pulse was followed by acquisition of a full plane k-space data using centric phase encoding.

#### MRI image analyses

The tumor volume was determined by manually tracing the tumor boundary on T2W images to generate tumor ROIs, summing the areas from all ROIs and multiplying the result by the slice thickness. DWI, DCE and MTR metrics were estimated over the same ROIs.

#### DW-MRI metrics

Radial k-space data were reconstructed to images using custom python codes (https://github.com/PennPancreaticCancerImagingResource/DWIProcessing). The reconstructed images were subject to pixel-wise least squares fit to Equation [1] using all five b-values as described previously (19), where *S_0_, ADC*, and *KI* are the fitting parameters: *S_0_* is the signal intensity in the absence of diffusion weighting and *KI* (kurtosis index) is a constant that accounts for contributions from water molecules undergoing slow hence non-gaussian diffusion to the observed signal *S(b)*.

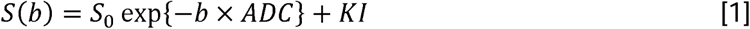

#### DCE-MRI metrics

For kinetic modeling of the DCE series, a reference region model (RRM) using muscle as the reference tissue was employed (19,28). The RRM was coded in custom Python code to derive the DCE-MRI metrics including the transfer constant of contrast agent from capillaries to interstitial space (*K^trans^* in /min) and extracellular/extravascular volume fraction (*V_e_*). *K^trans^* and *V_e_* of the reference tissue, the spinal muscle, were assumed to be 0.10/min and 0.10, respectively (19,28), and 4.6/s·mM was used as the relaxivity of the contrast media.

#### MTR metrics

MTR value was calculated pixel-wise by equation [2] to generate MTR maps from tumor ROIs:

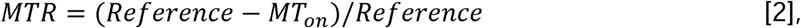

Where the *MT_on_* and *Reference* image were acquired with and without saturation of motion-restricted water, respectively. (19)

### Ex vivo studies

#### IHC Analyses

Upon completion of 2-week trial, tumors were harvested upon euthanasia. For IHC study, FFPE sections of the tumor were stained for H&E, CD31 and SR (one section for each stain per tumor) by the Pathological Core at CHOP as we described previously (19). The stained sections were scanned x40 using Aperio ScanScope CS2 and analyzed in QuPath by a GI pathologist (EEF) blinded to the treatment information. To avoid the bias inherent with selecting field of views for analysis, the entire section was analyzed using a pixel classifier method (19), where the tumor bed was segmented while lymph nodes and non-tumor areas (i.e., pancreas/chronic pancreatitis) were excluded from the analysis.

#### Single cell RNA sequencing (scRNAseq) and analyses

Tumors from mice in CNTRL, NGC, NGC+Cal treatment group were harvested upon euthanasia. Collection of viable cells encompassed a series of steps: tumor dissociation, lysis of red blood cells, clearance of debris, and dead cells removal. The cells then were suspended in 0.04% PBS in concentration exceeding 200K cells/100μL and were submitted to the Center for Applied Genomics core at Children’s Hospital of Philadelphia for scRNAseq. Next-generation sequencing libraries were prepared using the 10x Genomics Chromium Single Cell 3’ Reagent kit v3 per manufacturer’s instructions. Libraries are uniquely indexed using the Chromium dual Index Kit, pooled, and sequenced on an Illumina NovaSeq 6000 sequencer in a paired-end, dual indexing run.

Sequencing for each library was targeted at 20,000 reads per cell. Data was processed using the Cell Ranger pipeline (10x Genomics, v.6.1.2) for demultiplexing and alignment of sequencing reads to the mm10 transcriptome and creation of feature-barcode matrices.

Using the Cell Ranger (10X Genomics) output of filtered matrices supplied by the sequencing core, the Seurat package (v4) is used to integrate the data for each sample, followed by SCTransform (v2) to perform normalization and unsupervised clustering. In Bioconductor (v3.16), the SingleR package combined celldex databases is used to perform automated cell-type assignments. Cirrocumulus was used for interactive exploration and visualization of datasets. Heatmaps and violin plots were generated using Cirrocumulus. Target genes were extracted from literature (29–32).

## Statistical analysis

Statistical analyses were conducted either in GraphPad Prism (version 10.1.2, San Diego, CA) or using R statistical software (version 4.3.1), assuming an alpha level of 0.05.

For each mouse, tumor volume was measured by MRI (on T2W images) at baseline, on Day-7 and Day-14. As a measure of tumor progression, the percentage (%) change in tumor size between baseline and Day-14 was calculated by 100%*(tumor size at day14 - day0)/day0.

Statistical analyses were conducted in GraphPad or by R software with α value set at 0.05. For MRI data (tumor size, ADC, KI, Ktrans, Ve and MTR), except tumor size they all passed the normality test (Shapiro-Wilk). Two-tailed t-test and Mann-Whitney non-parametric test were applied. For IHC data, Mann-Whitney non-parametric test was applied to address the outlier bias. To assess the likelihood of tumors doubling in size or undergoing regression differs significantly in NGC+X (X =Cal, Losa) groups versus NGC alone, logistic regression analysis was applied with the outcome being whether the tumor size was doubled (Y/N) or was regressed (Y/N) while the main predictor being the treatment group.

## Results

### Inhibition of tumor growth and changes of cancer subtypes in response to chemotherapy alone and in combination with stromal intervention

Water-fall plots for tumor size change over the 2-week study period are shown in **Figure 2A-D** for individual mice enrolled in the 4 treatment arms while tumor size evolution is shown in **SI Figure 1**. Compared to the CNTRL group that exhibited 0% regression and majority of them doubled their size over the 14d-ay period (doubling rate = 89%), NGC mediated 22% regression rate with 25% doubling rate; NGC+Cal reduced the doubling rate to 14%, which was further reduced to 6% by NGC+Losa accompanied by 50% regression rate (**Figure 2E**). Tumor size changes comparing CNTRL vs all other treatments are highly significantly (*p* < 0.0001, **Figure 2F**). Notably, only NGC+LOSA but not NGC+CAL is more effective than NGC to inhibit tumor growth (*p* <0.05, **Figure 2F**). Statistical analysis suggested that NGC+Losa is capable of inhibiting more tumor growth than NGC (OR = 3.5, CI = 1.05-1.22, *p* < 0.05, **Figure 2G**) while NGC+Cal regime is not. As a marker of treatment mediated cell-kill, increased ADC was observed on Day-7 in NGC and NGC+Losa treated mice compared to the CNTRL (**SI Figure 1F**).

**Figure 2.**
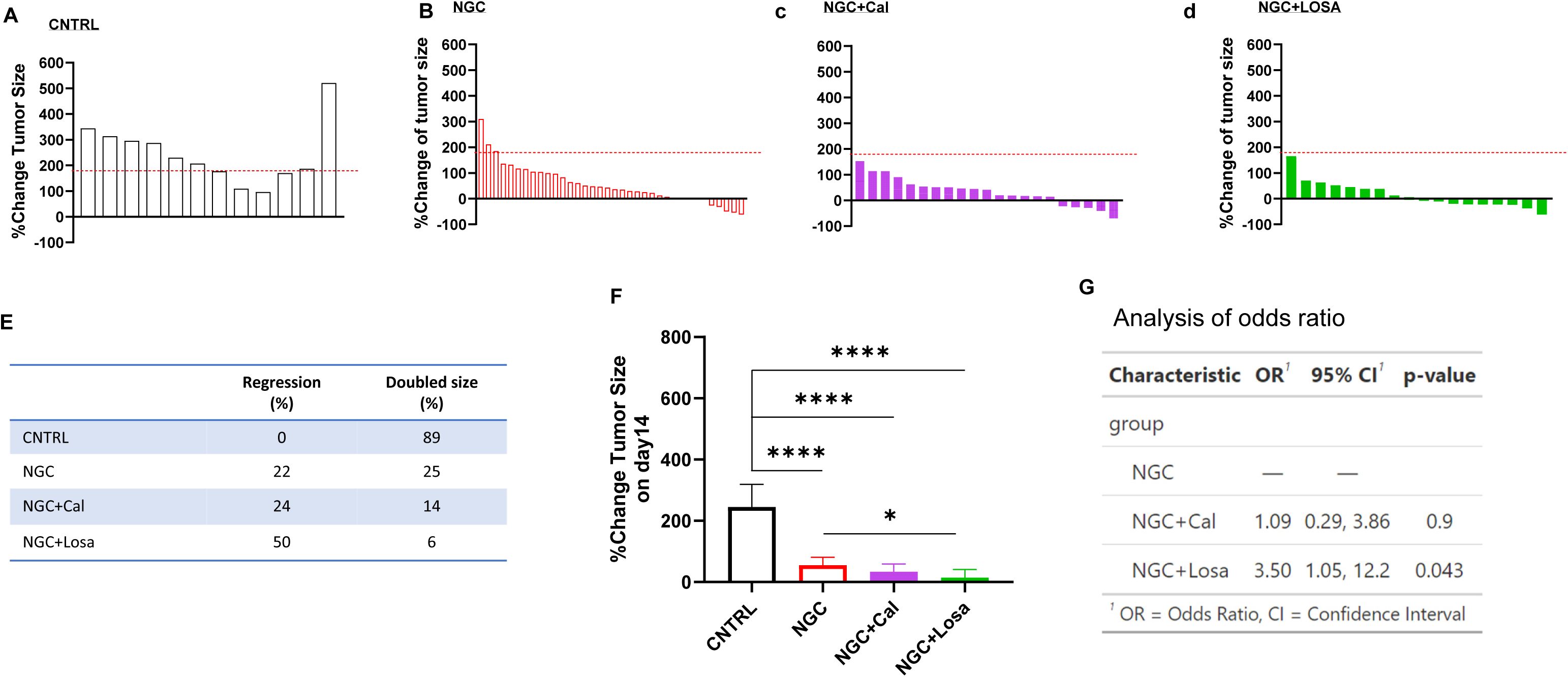
Water fall plots of tumor size change in response to stromal drug combined with chemotherapy in KPC model. Percent change of tumor size over the 14-day period was plotted for each KPC mouse enrolled in CNTRL (**A**), NGC (**B**), NGC+Cal (**C**), and NGC+Losa (**D**) group. Bars arising above the red dotted line are tumors which doubled their volume, whereas bars of negative values are tumors which regressed. Rate of regression and doubling size for each treatment group (**E**). Comparison of %tumor size change on Day-14. Odds ratio analysis to compare NGC+X (X= Cal, Losa) with NGC (**F**). **** *p* < 0.0001, * p < 0.05 by two tailed t-test.

UMAP derived from unsupervised clustering of all viable cells from four KPC tumors is shown in **SI Figure 2A**: Clusters were assigned as cancer cells (0 and 2) (30,33), fibroblast cells (8 and 10) (31) and other types of cells distributed differently corresponding to specified treatment (**SI Figure 2B-E)**. NGC or NGC+X (X= Cal, Losa) treatment reduced the %cancer cells in the tumor by a factor > 2 of CNTRL, with NGC+Losa being the most effective in eradicating cancer cells. Furthermore, NGC+Losa treatment led to dramatic increases of T and B lymphocyte infiltration (**SI Figure 2B-E,F**), consistent with the significantly enhanced inhibition of tumor growth over NGC alone (Figure 2F).

scRNAseq analyses revealed cancer subtype (E *versus* M) alteration by these treatments (**Figure 3A**): NGC treatment enriched the M subtype compared to CNTRL. Notably, NGC+Cal reversed completely the E/M ratio of NGC, and this appears unique to Cal. We applied a different method to query the data and arrived similar results (**SI Figure 3**). We then compared the transcriptome of cancer cells (cluster 0, 2) from different treatment groups against E and M panel genes defined in other scRNAseq studies of KPC model (29,30) (**SI Figure 4**). We found among E panel genes, Tff1 (Trefoil factor family 1) expression was reduced remarkably by NGC treatment (red arrow) whereas Fn1 (Fibronectin 1), a M panel signature, was notably increased by NGC (**Figure 3B**); NGC+Cal or NGC+Losa reversed the NGC-induced changes in Tff1 and Fn1 expression towards CNTRL level. Furthermore, gene set enrichment analysis (GSEA) of epithelial mesenchymal transition (EMT) - associated genes revealed a positive normalized enrichment score (NES) after NGC treatment corresponding to upregulation of EMT genes in contrast to negative NES after NGC+X (X = Cal, Losa) treatment, i.e., downregulation of EMT (**Figure 3C**).

**Figure 3.**
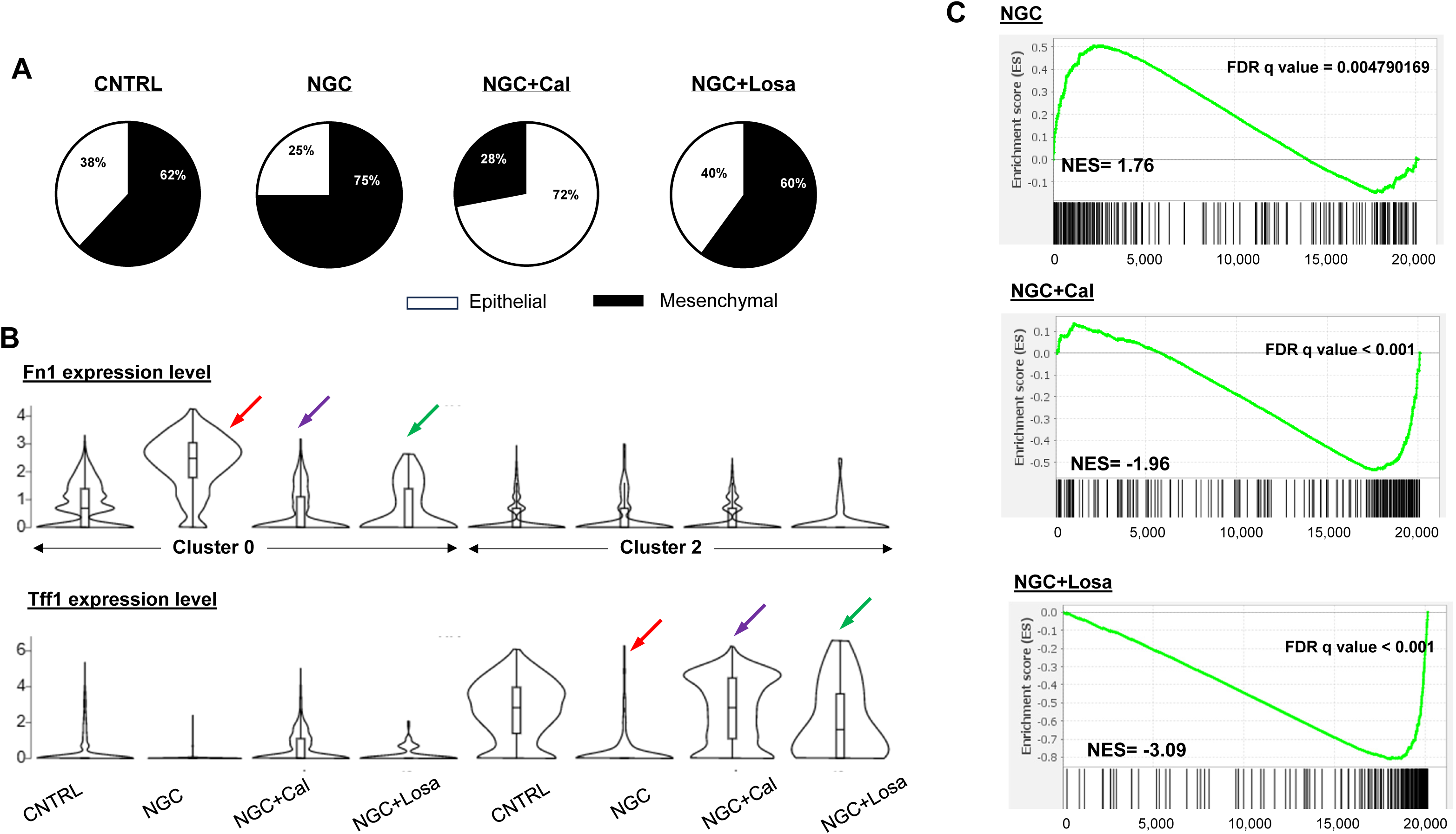
scRNAseq analysis revealed transcriptome changes after chemotherapy and in combination with calcipotriol or losartan. Fraction of cancer cells of epithelial (E) or mesenchymal or quasi- mesenchymal (M) subtype (**A**). Violin plots of M subtype signature gene Fn1 and E signature gene Tff1 show distinct expression among treated sample (**B**). GSEA revealed activation of EMT program by NGC treatment, but not when NGC was combined with calcipotriol or losartan (**C**). NES = normalized enrichment score.

### Changes of microvasculature, matrix collagen and FAP signature in response to chemotherapy alone and in combination with stromal intervention

#### Microvasculature density and perfusion /permeability

A metric derived from DCE, K^trans^ is the multiplication of capillary permeability and perfusion however under the flow-limited condition as in PDAC tumor, K^trans^ may primarily represent vascular permeability. Tumor K^trans^ maps (in pseudo-color) of a mouse treated by NGC at Day-0 and 7 revealed a large reduction in K^trans^ and exhibited a rim/core pattern with higher K^trans^ at the tumor periphery compared to the center (**Figure 3A**). Group wise, NGC induced a significant reduction in K^trans^ on Day-7 compared to baseline (** *p* < 0.01, paired t-test, **Figure 3A**), and the reduction of K^trans^ is corroborated with a significantly decreased MVD by CD31 staining (* *p* < 0.05, **Figure 3C,D,G**). While MVD after NGC+Losa was significantly lower than CNTRL tumors (** *p* < 0.01, **Figure 3C,F,G**), K^trans^ measured at Day-7 did not show significant difference, likely due to small sample size (n= 6-8 in Figure 3A,B) relative to a high degree of inter-tumor heterogeneity exhibited in KPC model. Limited sample size /time point for DCE is due to the requirement of two successful placements of tail vein catheter (at Day-0 and 7), which is challenging in C57/B6 mice.

Compared to minimal change in CNTRL (3%) and NGC group (-3%) (**SI Figure 5C**), consistent reductions of V_e_ (fraction of extravascular extracellular space, EES) were detected in all NGC+Cal treated mice (\*\*\**p* < 0.001, **SI Figure 5A**) as well as in NGC+Losa treated tumors (\**p* < 0.05, **SI Figure 5B**). (34) extravascular extracellular space was significantly higher in primary malignant tumors compared with nontumoral pancreatic tissue downstream (P = .018). disorganized collagen was reported after gem +nab-paclitaxel but it seems the doing of nab-paclitaxel.

#### Matrix collagen content

Compared to relatively organized and mild collagen deposition in the CNTRL tumor (**Figure 5A**), Sirius red (SR)-stained micrographs revealed a dense accumulation of collagen in between the tumor nests after chemotherapy (**Figure 5B**) and chemo stromal treatments (**Figure 5C,D**). Compared to the CNTRL, tumor collagen content was increased significantly after NGC and NGC+Losa (** *p* < 0.01, **Figure 5E**). Addition of Cal to NGC mitigated NGC-induced collagen increase (*p* = 0.0675 compared to CNTRL, **Figure 5E**), consistent with Cal’s ability to reduce fibrosis reported previously (13).

**Figure 4.**
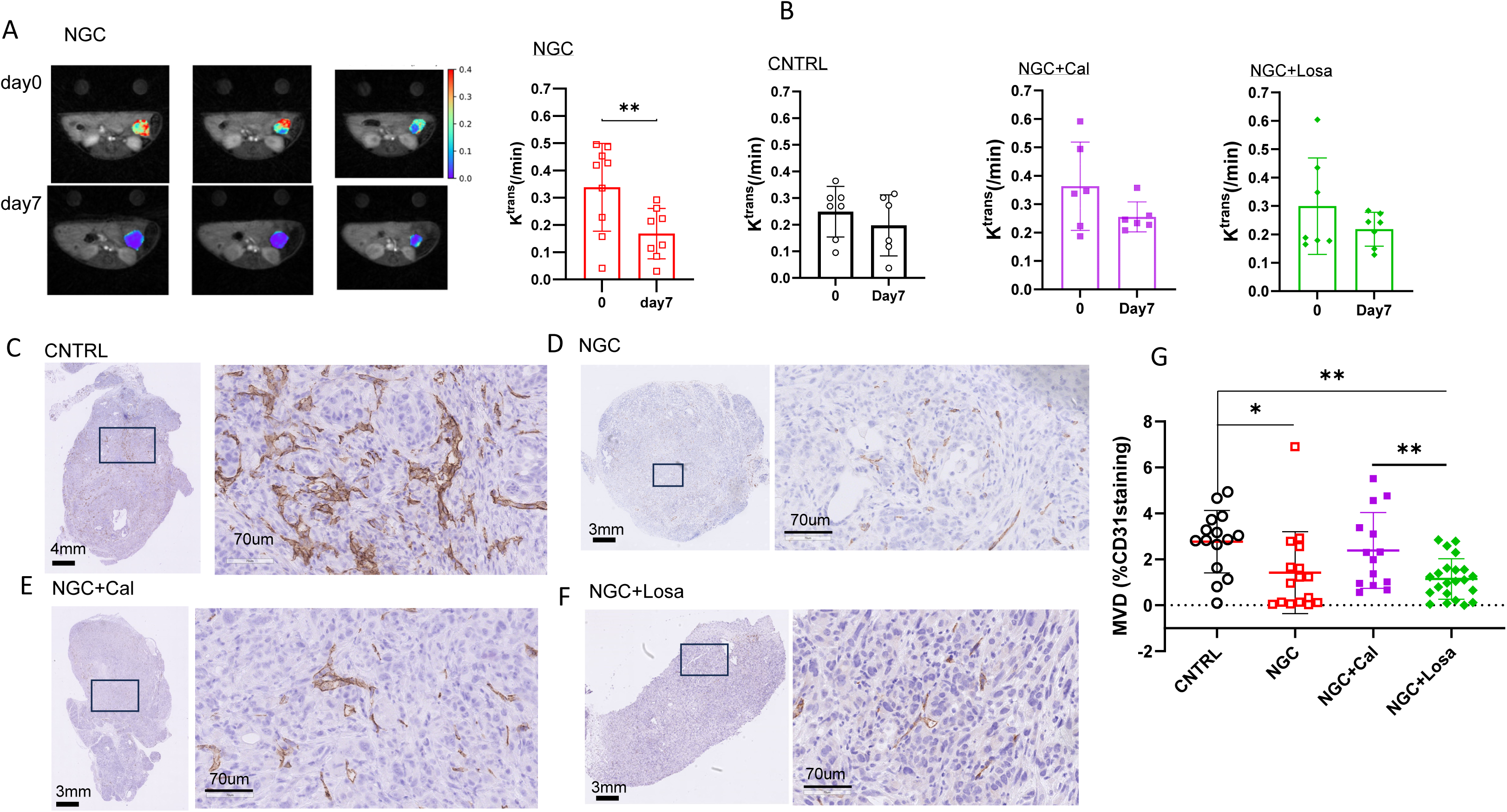
Treatment induced changes in DCE metric and MVD. Representative K^trans^ of the tumor from a mouse in NGC group and mean ± sem of K^trans^ of NGC group (**A**). Plots of from CNTRL, NGC+Cal and NGC+Losa (**B**). Representative CD31 stained sections from all treatment groups (**C-F**) and estimation of MVD based on CD31 staining (**G**).

**Figure 5.**
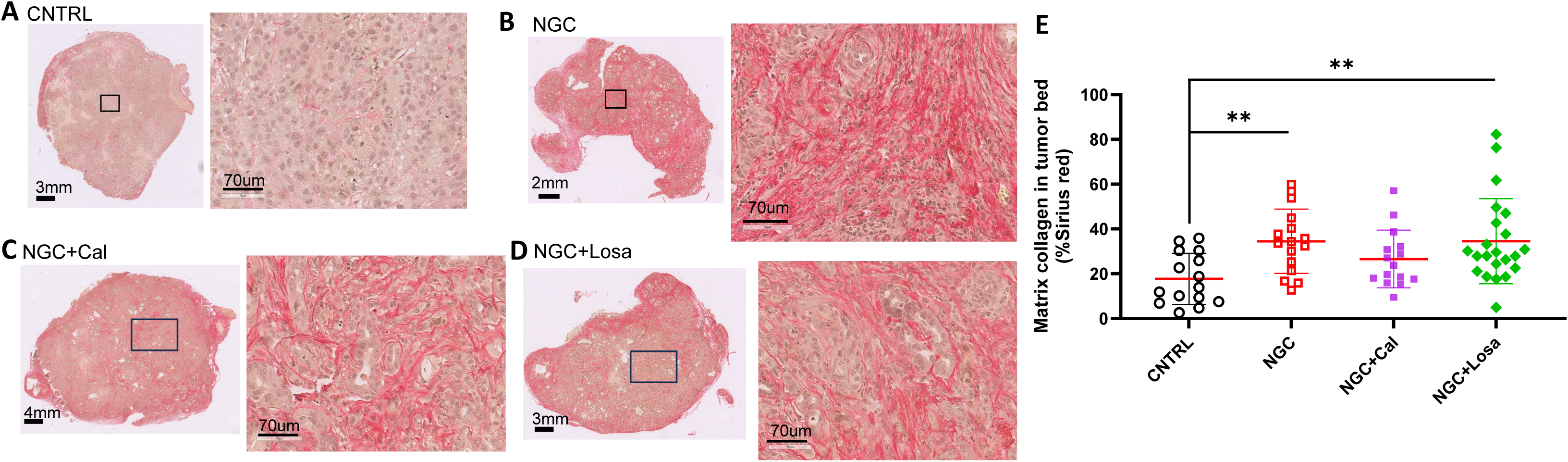
Treatment induced changes in collagen in KPC tumors assessed by IHC. Representative SR stained sections from all treatment groups (**A-D**). Collagen content was estimated by %SR staining in the tumor bed for all treatment groups (**E**). ** *p* < 0.01, * p < 0.05 by Mann-Whitney test.

#### FAP expression on CAFs

Compared to intense FAP staining in CNTRL tumors (**Figure 6A**), a remarkable clearance of FAP was observed in tumors after chemotherapy or chemo stromal therapy (**Figure 6B-D**). Highly significant reduction of FAP in NGC (\*\*\**p* < 0.001) and NGC+Losa (\*\**p* < 0.01) compared to CNTRL were demonstrated (**Figure 6E**). Reduced FAP levels in NGC and NGC+Losa treated tumors mirrored the increased tumor collagen after the same treatments (Figure 5E). Transcriptome analysis of CAFs (cluster 8, 10, **SI Figure 6**) revealed remarkably enhanced expression of iCAF related genes after NGC+Losa treatment (black arrow) and diminished expression of myCAF panel genes after NGC or NGC+stromal therapy (open arrow).

**Figure 6.**
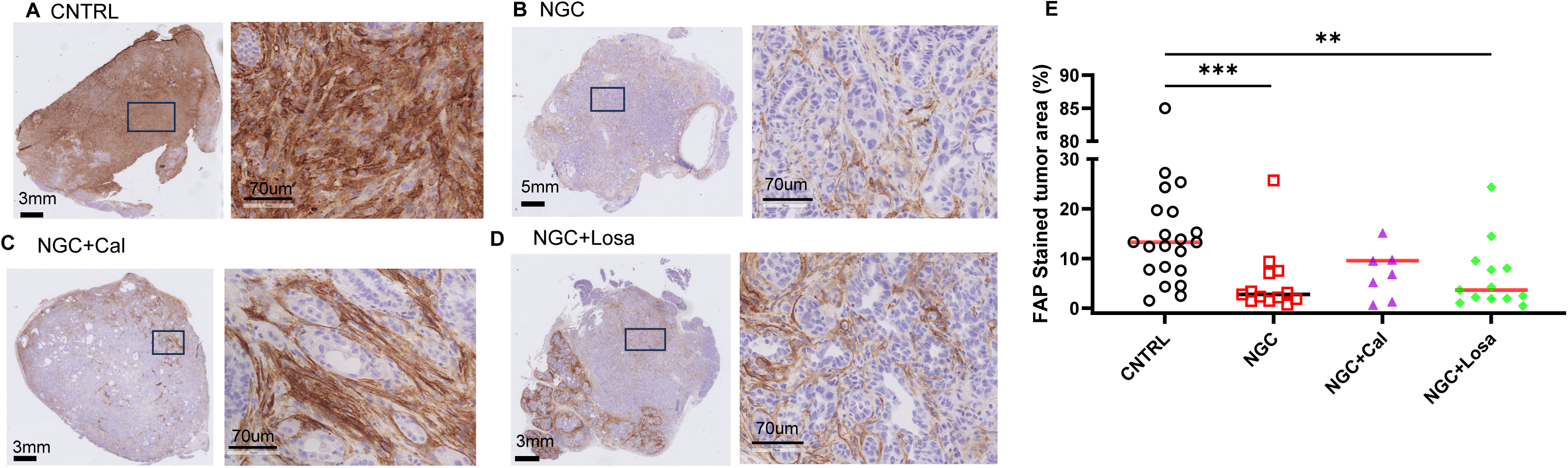
Treatment induced changes in FAP in KPC tumors assessed by IHC. Representative FAP stained sections from all treatment groups (**A-D**). FAP level was estimated by %positive staining in the tumor bed for all treatment groups (**E**). ** *p* < 0.01 by Mann-Whitney test.

## Discussion

In KPC model, which is relatively resistant to chemotherapy regimens, MRI-based tumor size was able to detect enhanced inhibition of tumor growth by chemo stromal therapy versus chemotherapy alone over a short trial period of 14 days. Early detection of microvascular function by DCE on Day-7 suggests that NGC chemotherapy significantly reduced microvascular permeability (K^trans^) as well as MVD (Figure 4), supporting the pruning effect of chemotherapy that reduces angiogenesis by killing proliferating cancer and endothelial cells (35). However, NGC+Cal (or Losa) did not reduce K^trans^ significantly, suggesting that stroma-directed agents might have an effect to increase K^trans^, especially for Cal, which is shown to reprogram PDAC stroma from inflammatory to quiescent phenotype (13) and NGC+Cal treatment did not affect MVD compared to CNTRL (Figure 4G). ADC was measured on Day-0 (baseline) and then 7 days after each chemotherapy administration. Compared to Day-0 and 7, CNTRL group exhibited a reduction of ADC in contrast to an increase of ADC in NGC+Losa group (SI Figure 1E), consistent with the biophysical principle of ADC, which measures the diffusion distance traveled by water molecules in extracellular space (ECS) (36). In ECS, intact cell membranes impose a boundary that stops free diffusion of water molecules, and as such a region of high cell density would allow a short diffusion distance (low ADC) compared to a region with sparse cell density. Consequently, tumor growth would reduce ADC meanwhile, treatment induced cell death would cause cells to lose the membrane integrity and allow diffusion, leading to increased ADC, and more cell death, the higher ADC. Hence the therapeutic efficacy based on percent ADC change (SI Figure 1E) would suggest NGC+Losa > NGC > NGC+Cal. Indeed, ADC values from NGC and NGC+Losa treatment group are significantly higher than those of CNTRL on Day-7 (SI Figure 1F). The lack of correlation between ADC versus tumor size over time for all treatment groups (SI Figure 1A-D) suggests that ADC is an orthogonal marker of ECS fraction, relating to cellular architecture and microenvironment of the tumor. Also derived from DWI, kurtosis index (KI) is a measure of diffusion heterogeneity (37,38). Our data show that diffusion heterogeneity increases with tumor growth in CNTRL and on Day-14, KI was higher in CNTRL than all treatment groups while reaching statistical significance with NGC group (SI Figure 8).

Our data suggest distinct mechanisms of stroma drug Cal and Losa to enhance NGC chemotherapy. NGC+Losa treatment led to a higher degree of tumor growth inhibition than NGC (Figure 2) along with robust lymphocytes infiltration and near eradication of cancer cells (SI Figure 2E,F), two features clearly distinct from CNTRL tumor - this data is consistent with recent clinical study that shown Losa combined with FOLFIRINOX and Chemoradiation reduced immune suppressing T regulatory cells and FOXP3^+^ (15). NGC treatment enriched M subtype while reducing E subtype of cancer cells, and activated the EMT program (Figure 3) – this ex vivo observation was in line with earlier in vitro data that exposure of human PDA cell lines to FOLFIRINOX treatment led to enrichment of quasi mesenchymal state (30); interestingly, NGC+Cal reversed this trend and hindered the EMT program (Figure 3). Since enrichment of M subtype and EMT activation are known to drive chemoresistance in PDA (39), our data suggests that adding Cal to NGC may better maintain the chemosensitivity of PADC than adding Losa. Overall, the scRNAseq study although preliminary in nature due to limited sample size (n=1 for each treatment), provides plausible mechanism for enhanced therapeutic efficacy of chemo stromal therapy over chemotherapy alone.

NGC treatment induced a significant increase of stromal collagen content, consistent with tumor restricting effect of type-1 collagen opposing CAF’s tumor promoting role (40). The inverse relationship between SR and FAP staining level (high SR /low FAP and vice versa) revealed by our data suggest that FAP could be a marker for chemotherapy response since high FAP expression is correlated with poor prognosis /invasion / large tumor size (41). As a universal marker of CAFs, genetic deletion of FAP appears to delay tumor growth and leads to increased tumor collagen content and reduced myofibroblast content in KRAS^G12D^ - driven lung cancer (42) – these are consistent with our observations in KPC tumors (also carrying KRAS^G12D^ mutation) that treatment-induced FAP reduction is associated with increased collagen (Figure 5) and a trend of downregulation of myCAF related genes (open arrow in SI Figure 7).

Consistent with SR staining (Figure 5), tumor MTR in NGC group exhibited a trend of increase over time and the NGC+Losa group had the greatest MTR value on Day-14 compared to other groups (SI Figure 8). However, no significant correlation was observed between MTR and SR staining (R^2^ = 0.11, *p* = 0.25), although we have limited cases which have both Day-14 MTR and SR staining (n= 3, 3, 5, 2 for CNTRL, NGC, NGC+Cal and NGC+Losa, respectively).

However, it has been reported that MTR value could be dominated by changes in cellularity in highly proliferative malignancy (24) or in response to therapies that induce a high level of cell death leading to releasing of intracellular bound water (21).

In conclusion, in a highly clinically relevant GEM model of PDAC, we have shown the efficacy of standard care chemotherapy (NGC) can be significantly enhanced by combination with stromal agent, Cal (a synthetic vitamin D) or Losa (a hypertension drug). DCE and CD31 staining suggest that NGC chemotherapy reduced the vascular permeability and MVD, providing a physical mechanism for chemoresistance, while the enrichment of M/QM subtype cancer cells revealed a molecular mechanism. Addition of Losa to NGC appears to enhance lymphocyte infiltration dramatically. Meanwhile, addition of Cal to NGC reversed chemotherapy- induced enrichment of M/QM subtype. From these preliminary scRNAseq study, we generated useful hypotheses which can be tested in future mechanistic studies.

## Acknowledgement

The studies are partially supported by Co-clinical Imaging Resource Program (CIRP) of NCI (U24CA231858). We appreciate the institutional resources including the Mouse Hospital of the Pancreatic Cancer Research Center (PCRC), University of Pennsylvania, and Small Animal Imaging Facility (SAIF), Department of Radiology, University of Pennsylvania.

---

**SI Figure 1.**
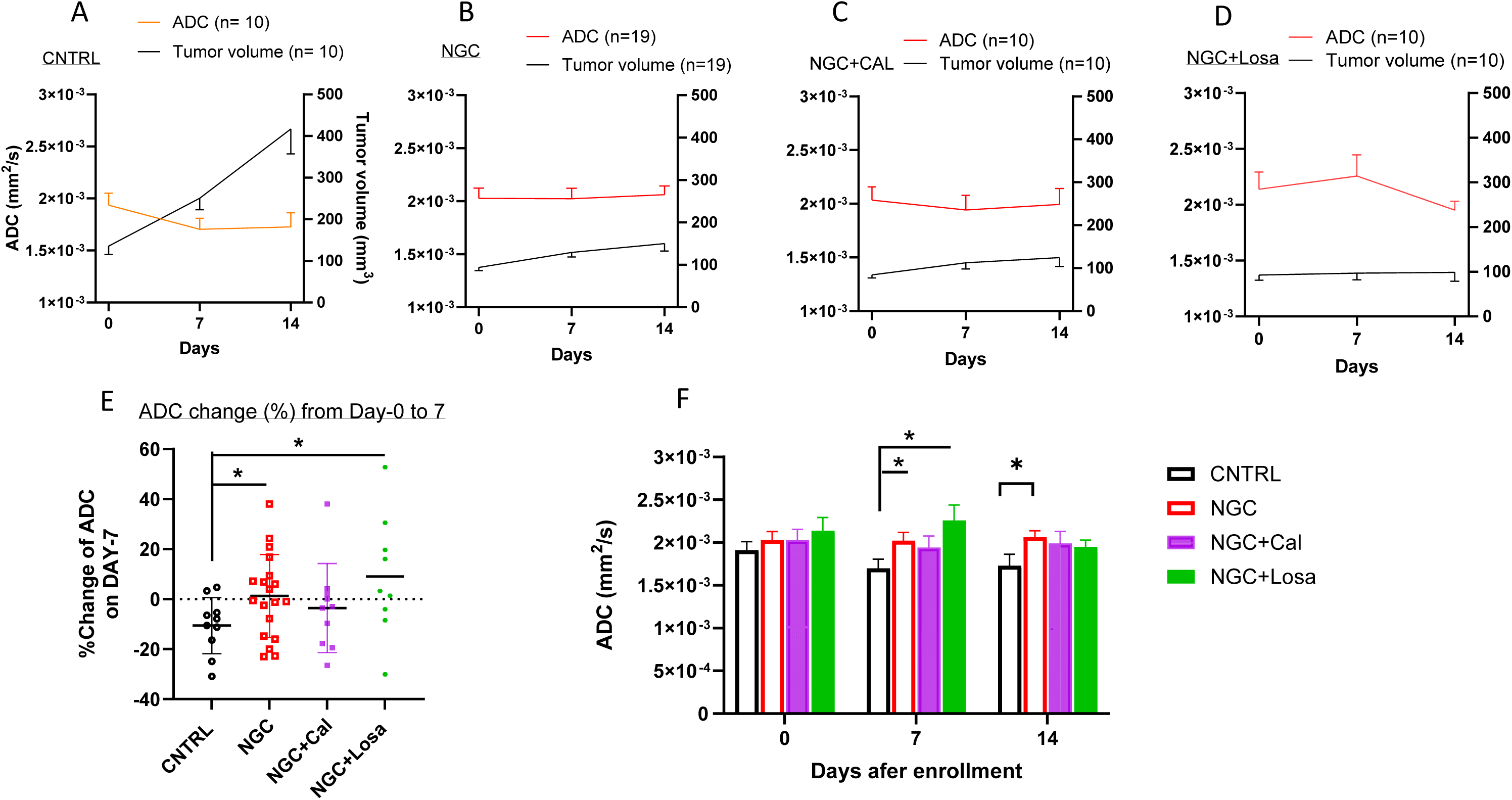
ADC and tumor size time course for all treatment groups. ADC and tumor size were plotted for all mice which underwent DWI in each treatment group (**A-D**). Percent change of ADC on Day-7 (**E**). Comparison of group ADC (mean ± SD) for all groups on Day-0, 7 and 14 (**F**). Only negative or positive part of the error bar (standard deviation) are shown for clarity. * p < 0.05 by two-tailed t-test.

**SI Figure 2.**
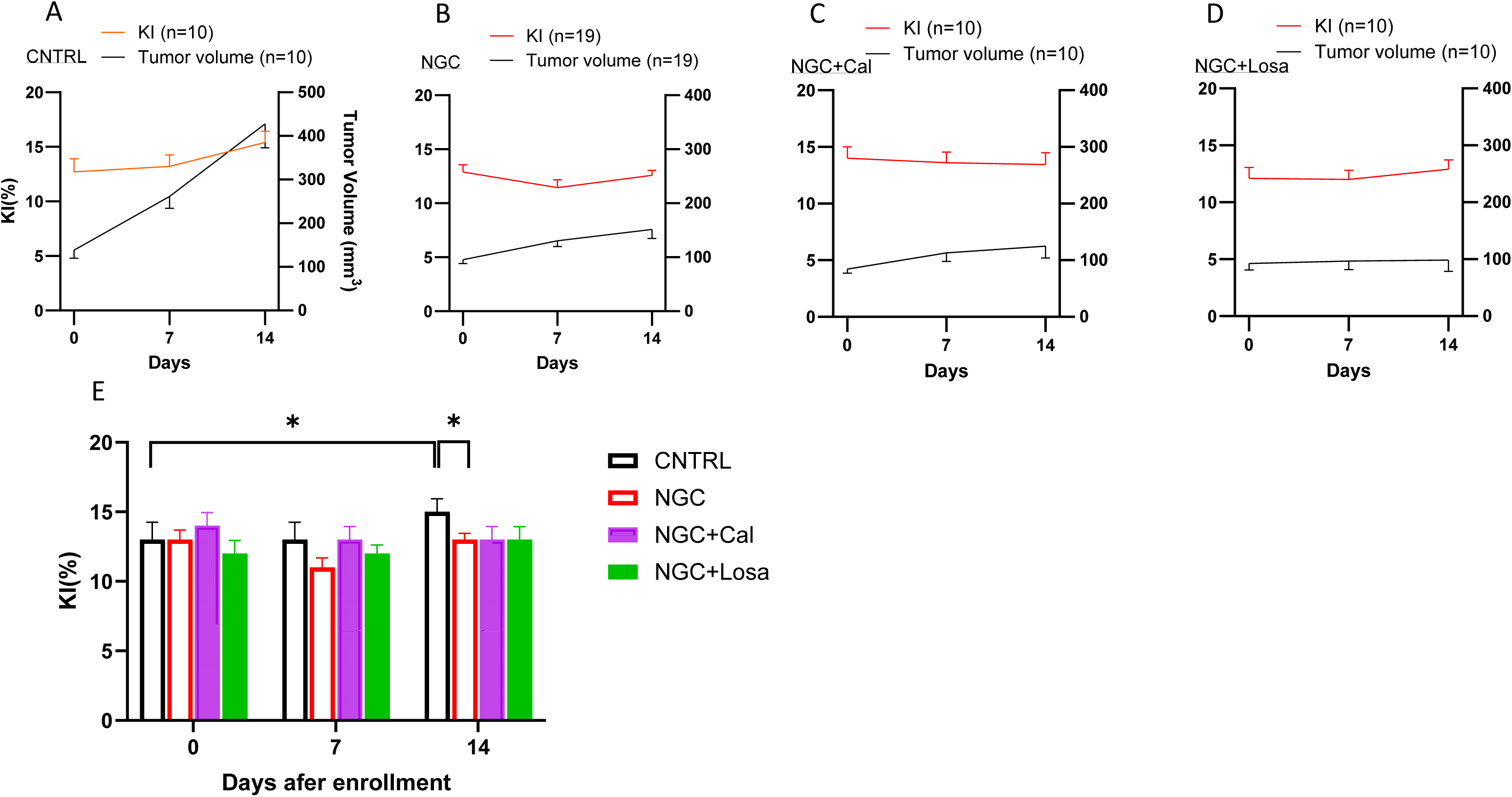
KI and tumor size time course for all treatment groups. KI and tumor size were plotted for all mice which underwent DWI in each treatment group (**A-D**). Comparison of group KI (mean ± SD) for all groups on Day-0, 7 and 14 (**E**). * p < 0.05 by two-tailed t-test.

**SI Figure 3.**
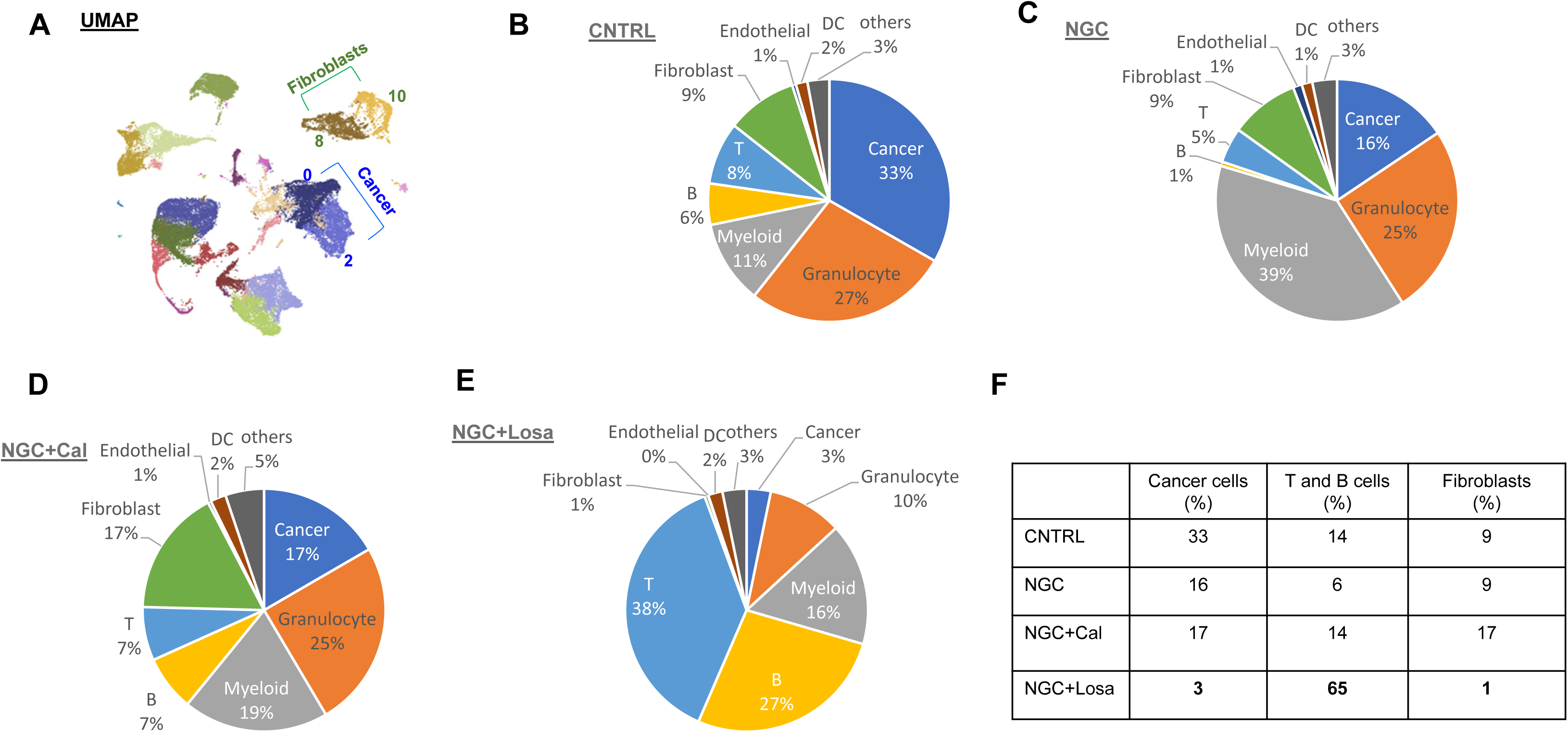
Uniform Manifold Approximation and Projection (UMAP) from unsupervised clustering and cell type distribution in tumor samples. UMAP of all samples (**A**). Cell types and distribution for CNTRL (**B**), NGC (**C**), NGC+Cal (**D**), NGC+Losa (**E**) (n =1 for each treatment). Comparison of percentage of cancer, T cell & B cell and fibroblast cell populations after each treatment (**F**).

**SI Figure 4.**
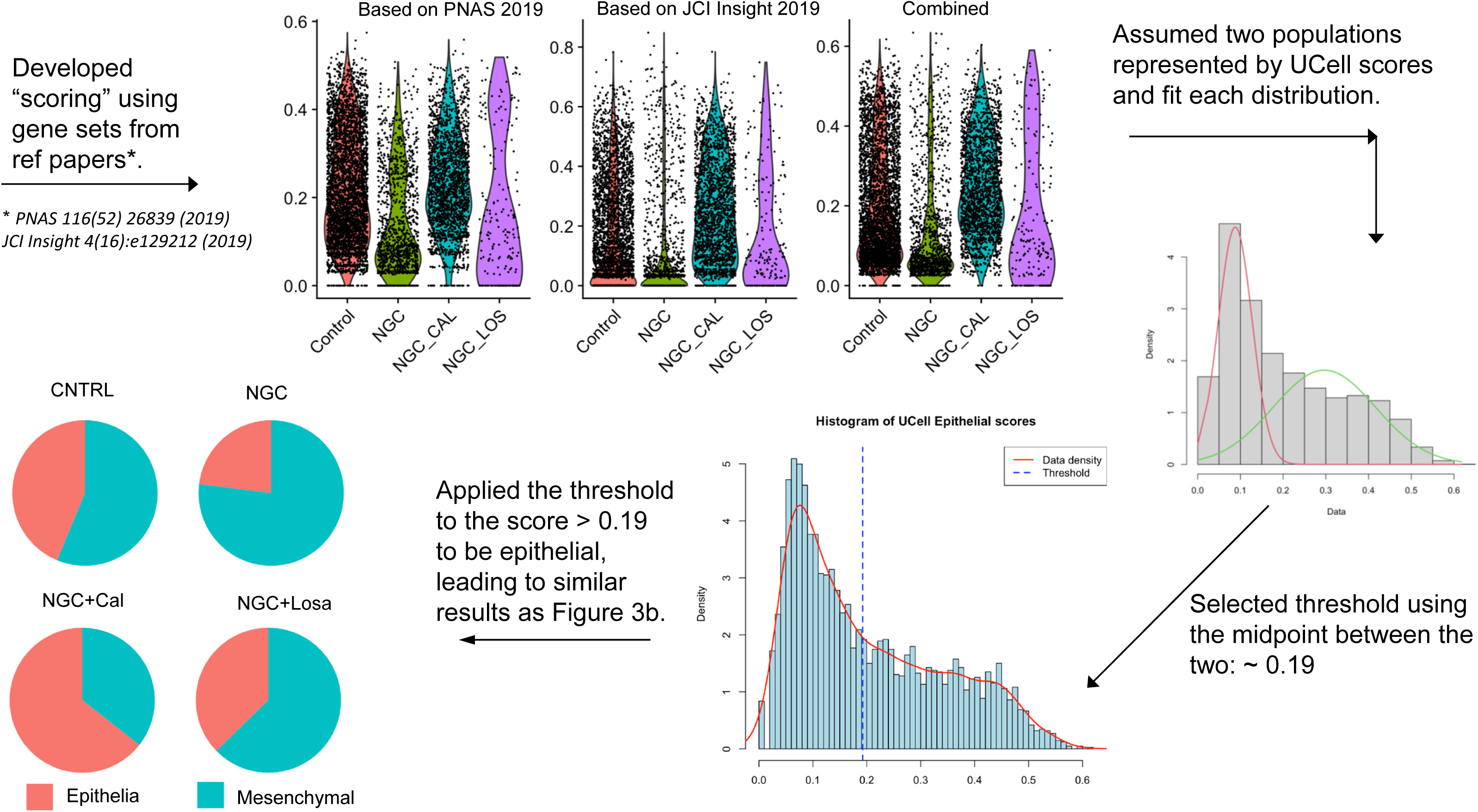
A different analytical approach to confirm Figure 3A results. A scoring using gene sets from ref papers was applied to cancer cell clusters of each sample; assuming two populations (E vs. M) represented by UCell scores, from which each distribution was fit and thresholded, leading to similar results as Figure 3A.

**SI Figure 5.**
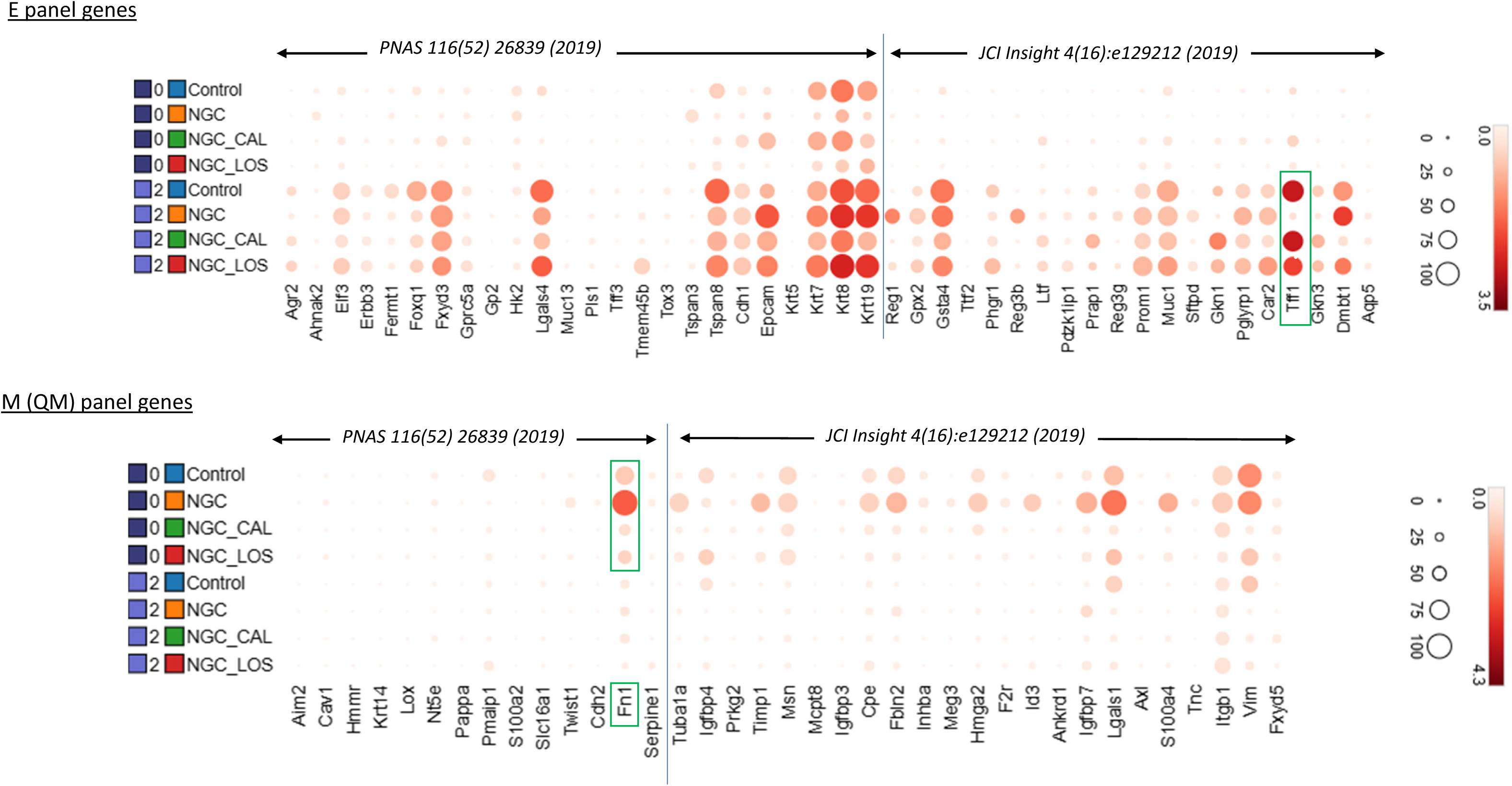
Transcriptome changes of cancer cells (cluster 0, 2) in E (epithelial) and M (mesenchymal or quasi mesenchymal) panel genes in CNTRL, NGC, NGC combined with Cal or Losa, respectively. Panel genes are based on references cited.

**SI Figure 6.**
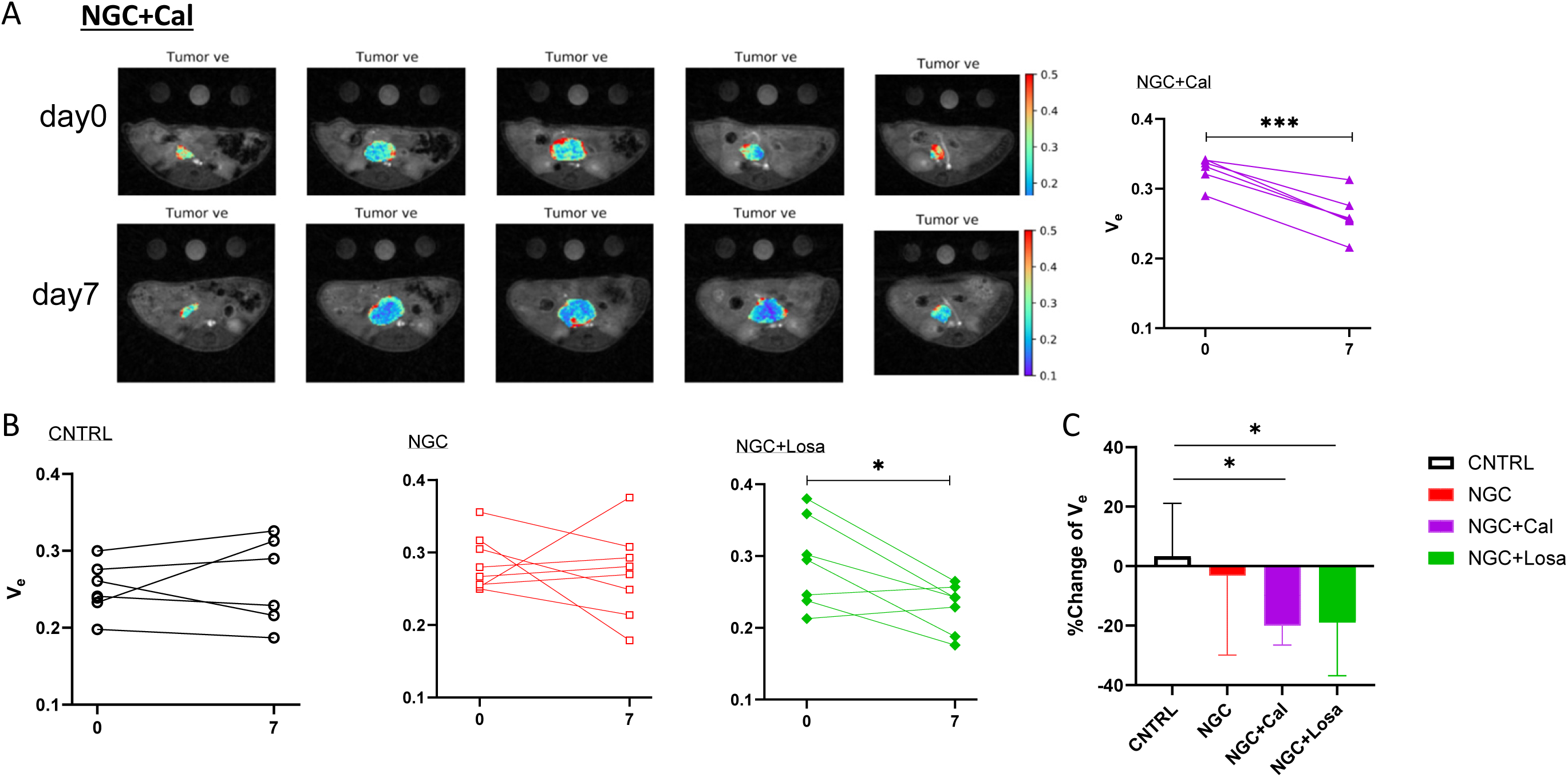
Treatment induced changes in DCE metric V_e_. Representative V_e_ map of the tumor from a mouse in NGC+Cal group and V_e_ on Day-0 and 7 (**A**). Five images from the same tumor are shown for Day-0 and 7, respectively. Plots of V_e_ from CNTRL, NGC and NGC+Losa (**B**). Change (%) of V_e_ for all treatment groups (**C**). \**P* <0.05, two-tailed t-test.

**SI Figure 7.**
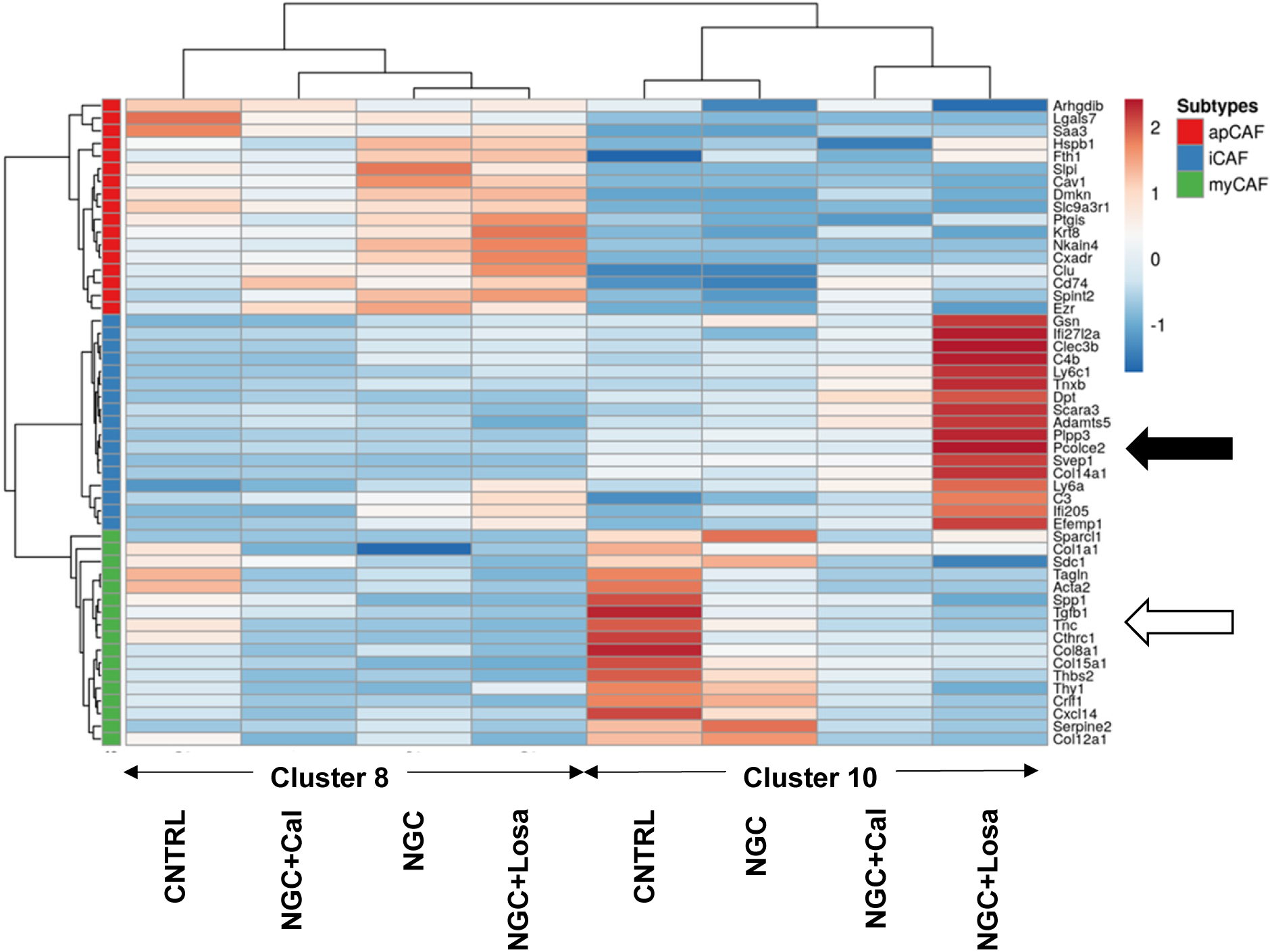
scRNAseq analysis revealed transcriptome changes in CAFs after chemotherapy and in combination with calcipotriol or losartan. CAFs are represented by cluster 8 and 10 and panel gens for apCAF, iCAF and myCAF were referenced to (32). Closed black arrow points to enhanced expression of iCAF panel genes after NGC+Losa treatment. Open arrow points to reduced expression of myCAF in treated tumors compared to CNTRL.

**SI Figure 8.**
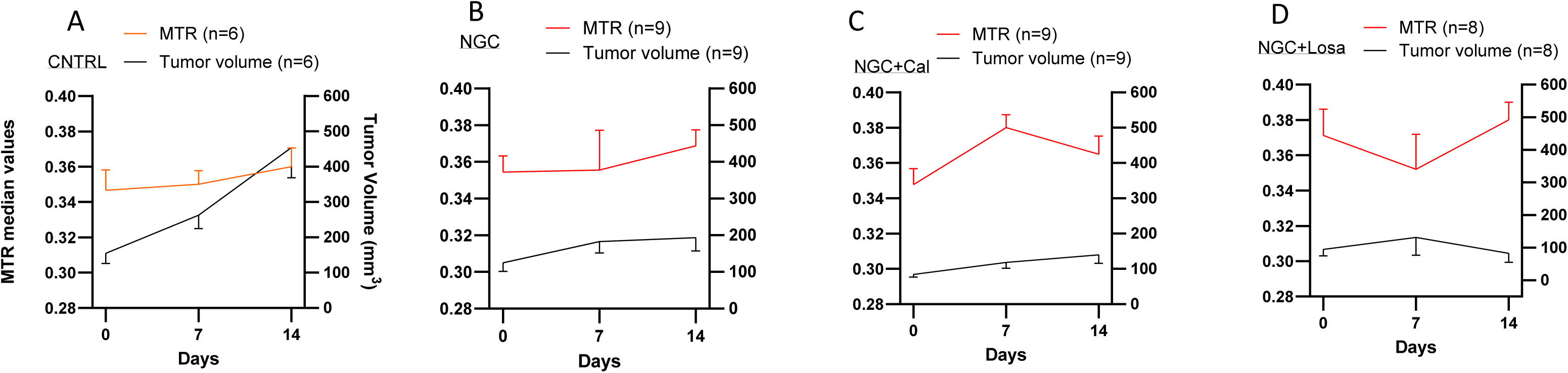
MTR and tumor size time course for all treatment groups. MTR and tumor size were plotted for all mice which underwent MTR study in each treatment group (**A-D**).

## Notes

### Competing Interest Statement

The authors have declared no competing interest.

## References

1. Lee HS, Park SW. Systemic Chemotherapy in Advanced Pancreatic Cancer. Gut and liver 2016;10(3):340–347.

2. Springfeld C, Jäger D, Büchler MW, Strobel O, Hackert T, Palmer DH, Neoptolemos JP. Chemotherapy for pancreatic cancer. Presse medicale (Paris, France : 1983) 2019;48(3 Pt 2):e159–e174.

3. Von Hoff DD, Ervin T, Arena FP, Chiorean EG, Infante J, Moore M, Seay T, Tjulandin SA, Ma WW, Saleh MN, Harris M, Reni M, Dowden S, Laheru D, Bahary N, Ramanathan RK, Tabernero J, Hidalgo M, Goldstein D, Van Cutsem E, Wei X, Iglesias J, Renschler MF. Increased Survival in Pancreatic Cancer with nab-Paclitaxel plus Gemcitabine. New England Journal of Medicine 2013;369(18):1691–1703.

4. Ahmad SA, Duong M, Sohal DPS, Gandhi NS, Beg MS, Wang-Gillam A, Wade JL, 3rd,Chiorean EG, Guthrie KA, Lowy AM, Philip PA, Hochster HS. Surgical Outcome Results From SWOG S1505: A Randomized Clinical Trial of mFOLFIRINOX Versus Gemcitabine/Nab-paclitaxel for Perioperative Treatment of Resectable Pancreatic Ductal Adenocarcinoma. Ann Surg 2020;272(3):481–486.

5. Moffitt RA, Marayati R, Flate EL, Volmar KE, Loeza SG, Hoadley KA, Rashid NU, Williams LA, Eaton SC, Chung AH, Smyla JK, Anderson JM, Kim HJ, Bentrem DJ, Talamonti MS, Iacobuzio-Donahue CA, Hollingsworth MA, Yeh JJ. Virtual microdissection identifies distinct tumor- and stroma-specific subtypes of pancreatic ductal adenocarcinoma. Nature genetics 2015;47(10):1168–1178.

6. Bailey P, Chang DK, Nones K, Johns AL, Patch A-M, Gingras M-C, Miller DK, Christ AN, Bruxner TJC, Quinn MC, Nourse C, Murtaugh LC, Harliwong I, Idrisoglu S, Manning S, Nourbakhsh E, Wani S, Fink L, Holmes O, Chin V, Anderson MJ, Kazakoff S, Leonard C, Newell F, Waddell N, Wood S, Xu Q, Wilson PJ, Cloonan N, Kassahn KS, Taylor D, Quek K, Robertson A, Pantano L, Mincarelli L, Sanchez LN, Evers L, Wu J, Pinese M, Cowley MJ, Jones MD, Colvin EK, Nagrial AM, Humphrey ES, Chantrill LA, Mawson A, Humphris J, Chou A, Pajic M, Scarlett CJ, Pinho AV, Giry-Laterriere M, Rooman I, Samra JS, Kench JG, Lovell JA, Merrett ND, Toon CW, Epari K, Nguyen NQ, Barbour A, Zeps N, Moran-Jones K, Jamieson NB, Graham JS, Duthie F, Oien K, Hair J, Grützmann R, Maitra A, Iacobuzio-Donahue CA, Wolfgang CL, Morgan RA, Lawlor RT, Corbo V, Bassi C, Rusev B, Capelli P, Salvia R, Tortora G, Mukhopadhyay D, Petersen GM, Munzy DM, Fisher WE, Karim SA, Eshleman JR, Hruban RH, Pilarsky C, Morton JP, Sansom OJ, Scarpa A, Musgrove EA, Bailey U-MH, Hofmann O, Sutherland RL, Wheeler DA, Gill AJ, Gibbs RA, Pearson JV, Waddell N, Biankin AV, Grimmond SM, Australian Pancreatic Cancer Genome I. Genomic analyses identify molecular subtypes of pancreatic cancer. Nature 2016;531(7592):47-52.

7. Collisson EA, Sadanandam A, Olson P, Gibb WJ, Truitt M, Gu S, Cooc J, Weinkle J, Kim GE, Jakkula L, Feiler HS, Ko AH, Olshen AB, Danenberg KL, Tempero MA, Spellman PT, Hanahan D, Gray JW. Subtypes of pancreatic ductal adenocarcinoma and their differing responses to therapy. Nat Med 2011;17(4):500–503.

8. Dreyer SB, Chang DK, Bailey P, Biankin AV. Pancreatic Cancer Genomes: Implications for Clinical Management and Therapeutic Development. Clin Cancer Res 2017;23(7):1638–1646.

9. Collisson EA, Bailey P, Chang DK, Biankin AV. Molecular subtypes of pancreatic cancer. Nature Reviews Gastroenterology & Hepatology 2019;16(4):207–220.

10. Valkenburg KC, de Groot AE, Pienta KJ. Targeting the tumour stroma to improve cancer therapy. Nature reviews Clinical oncology 2018;15(6):366–381.

11. Provenzano PP, Cuevas C, Chang AE, Goel VK, Von Hoff DD, Hingorani SR. Enzymatic targeting of the stroma ablates physical barriers to treatment of pancreatic ductal adenocarcinoma. Cancer Cell 2012;21(3):418–429.

12. Miyamoto H, Murakami T, Tsuchida K, Sugino H, Miyake H, Tashiro S. Tumor-stroma interaction of human pancreatic cancer: acquired resistance to anticancer drugs and proliferation regulation is dependent on extracellular matrix proteins. Pancreas 2004;28.

13. Sherman Mara H, Yu Ruth T, Engle Dannielle D, Ding N, Atkins Annette R, Tiriac H, Collisson Eric A, Connor F, Van Dyke T, Kozlov S, Martin P, Tseng Tiffany W, Dawson David W, Donahue Timothy R, Masamune A, Shimosegawa T, Apte Minoti V, Wilson Jeremy S, Ng B, Lau Sue L, Gunton Jenny E, Wahl Geoffrey M, Hunter T, Drebin Jeffrey A, O’Dwyer Peter J, Liddle C, Tuveson David A, Downes M, Evans Ronald M. Vitamin D Receptor-Mediated Stromal Reprogramming Suppresses Pancreatitis and Enhances Pancreatic Cancer Therapy. Cell 2014;159(1):80–93.

14. Murphy JE, Wo JY, Ryan DP, Clark JW, Jiang W, Yeap BY, Drapek LC, Ly L, Baglini CV, Blaszkowsky LS, Ferrone CR, Parikh AR, Weekes CD, Nipp RD, Kwak EL, Allen JN, Corcoran RB, Ting DT, Faris JE, Zhu AX, Goyal L, Berger DL, Qadan M, Lillemoe KD, Talele N, Jain RK, DeLaney TF, Duda DG, Boucher Y, Fernández-Del Castillo C, Hong TS. Total Neoadjuvant Therapy With FOLFIRINOX in Combination With Losartan Followed by Chemoradiotherapy for Locally Advanced Pancreatic Cancer: A Phase 2 Clinical Trial. JAMA oncology 2019;5(7):1020–1027.

15. Boucher Y, Posada JM, Subudhi S, Kumar AS, Rosario SR, Gu L, Kumra H, Mino-Kenudson M, Talele NP, Duda DG, Fukumura D, Wo JY, Clark JW, Ryan DP, Fernandez-Del Castillo C, Hong TS, Pittet MJ, Jain RK. Addition of Losartan to FOLFIRINOX and Chemoradiation Reduces Immunosuppression-Associated Genes, Tregs, and FOXP3+ Cancer Cells in Locally Advanced Pancreatic Cancer. Clin Cancer Res 2023;29(8):1605–1619.

16. Schwartz LH, Litière S, de Vries E, Ford R, Gwyther S, Mandrekar S, Shankar L, Bogaerts J, Chen A, Dancey J, Hayes W, Hodi FS, Hoekstra OS, Huang EP, Lin N, Liu Y, Therasse P, Wolchok JD, Seymour L. RECIST 1.1-Update and clarification: From the RECIST committee. European journal of cancer (Oxford, England : 1990) 2016; 62:132-137.

17. Hosein AN, Brekken RA, Maitra A. Pancreatic cancer stroma: an update on therapeutic targeting strategies. Nature reviews Gastroenterology & hepatology 2020;17(8):487–505.

18. Cao J, Pickup S, Clendenin C, Blouw B, Choi H, Kang D, Rosen M, O’Dwyer PJ, Zhou R. Dynamic Contrast-enhanced MRI Detects Responses to Stroma-directed Therapy in Mouse Models of Pancreatic Ductal Adenocarcinoma. Clin Cancer Res 2019;25(7):2314–2322.

19. Romanello Joaquim M, Furth EE, Fan Y, Song HK, Pickup S, Cao J, Choi H, Gupta M, Cao Q, Shinohara R, McMenamin D, Clendenin C, Karasic TB, Duda J, Gee JC, O’Dwyer PJ, Rosen MA, Zhou R. DWI Metrics Differentiating Benign Intraductal Papillary Mucinous Neoplasms from Invasive Pancreatic Cancer: A Study in GEM Models. Cancers 2022;14(16):4017.

20. Kinh Do R, Reyngold M, Paudyal R, Oh JH, Konar AS, LoCastro E, Goodman KA, Shukla-Dave A. Diffusion-Weighted and Dynamic Contrast-Enhanced MRI Derived Imaging Metrics for Stereotactic Body Radiotherapy of Pancreatic Ductal Adenocarcinoma: Preliminary Findings. Tomography : a journal for imaging research 2020;6(2):261–271.

21. Gupta M, Choi H, Kemp SB, Furth EE, Pickup S, Clendenin C, Orlen M, Rosen M, Liu F, Cao Q, Stanger BZ, Zhou R. Quantitative MRI Measurements Capture Pancreatic Cancer and Stroma Reactions to New KRAS Inhibitor. bioRxiv 2024:2024.2011.2022.624844.

22. Jiang K, Ferguson CM, Ebrahimi B, Tang H, Kline TL, Burningham TA, Mishra PK, Grande JP, Macura SI, Lerman LO. Noninvasive Assessment of Renal Fibrosis with Magnetization Transfer MR Imaging: Validation and Evaluation in Murine Renal Artery Stenosis. Radiology 2017;283(1):77–86.

23. Li W, Zhang Z, Nicolai J, Yang G-Y, Omary RA, Larson AC. Magnetization transfer MRI in pancreatic cancer xenograft models. Magnetic Resonance in Medicine 2012;68(4):1291–1297.

24. Robison TH, Solipuram M, Heist K, Amouzandeh G, Lee WY, Humphries BA, Buschhaus JM, Bevoor A, Zhang A, Luker KE, Pettit K, Talpaz M, Malyarenko D, Chenevert TL, Ross BD, Luker GD. Multiparametric MRI to quantify disease and treatment response in mice with myeloproliferative neoplasms. JCI Insight 2022;7(19).

25. Kawase T, Yasui Y, Nishina S, Hara Y, Yanatori I, Tomiyama Y, Nakashima Y, Yoshida K, Kishi F, Nakamura M, Hino K. Fibroblast activation protein-α-expressing fibroblasts promote the progression of pancreatic ductal adenocarcinoma. BMC gastroenterology 2015;15:109.

26. Desai NP, Chunlin Tao C, Yang A, Louie L, Zheng T, Yao Z, Soon-Shiong P, Magdassi S. Protein stabilized pharmacologically active agents, methods for the preparation thereof and methods for the use thereof. US Patent 5,916,596A 1999.

27. Cao J, Song HK, Yang H, Castillo V, Chen J, Clendenin C, Rosen M, Zhou R, Pickup S. Respiratory Motion Mitigation and Repeatability of Two Diffusion-Weighted MRI Methods Applied to a Murine Model of Spontaneous Pancreatic Cancer. Tomography : a journal for imaging research 2021;7(1):66–79.

28. Pickup S, Romanello M, Gupta M, Song HK, Zhou R. Dynamic Contrast-Enhanced MRI in the Abdomen of Mice with High Temporal and Spatial Resolution Using Stack-of-Stars Sampling and KWIC Reconstruction. Tomography : a journal for imaging research 2022;8(5):2113–2128.

29. Hosein AN, Huang H, Wang Z, Parmar K, Du W, Huang J, Maitra A, Olson E, Verma U, Brekken RA. Cellular heterogeneity during mouse pancreatic ductal adenocarcinoma progression at single-cell resolution. JCI Insight 2019;4(16):e129212.

30. Porter RL, Magnus NKC, Thapar V, Morris R, Szabolcs A, Neyaz A, Kulkarni AS, Tai E, Chougule A, Hillis A, Golczer G, Guo H, Yamada T, Kurokawa T, Yashaswini C, Ligorio M, Vo KD, Nieman L, Liss AS, Deshpande V, Lawrence MS, Maheswaran S, Fernandez-Del Castillo C, Hong TS, Ryan DP, O’Dwyer PJ, Drebin JA, Ferrone CR, Haber DA, Ting DT. Epithelial to mesenchymal plasticity and differential response to therapies in pancreatic ductal adenocarcinoma. Proceedings of the National Academy of Sciences 2019;116(52):26835–26845.

31. Chen Y, Kim J, Yang S, Wang H, Wu CJ, Sugimoto H, LeBleu VS, Kalluri R. Type I collagen deletion in αSMA(+) myofibroblasts augments immune suppression and accelerates progression of pancreatic cancer. Cancer Cell 2021;39(4):548–565.e546.

32. Elyada E, Bolisetty M, Laise P, Flynn WF, Courtois ET, Burkhart RA, Teinor JA, Belleau P, Biffi G, Lucito MS, Sivajothi S, Armstrong TD, Engle DD, Yu KH, Hao Y, Wolfgang CL, Park Y, Preall J, Jaffee EM, Califano A, Robson P, Tuveson DA. Cross-Species Single-Cell Analysis of Pancreatic Ductal Adenocarcinoma Reveals Antigen-Presenting Cancer-Associated Fibroblasts. Cancer Discov 2019;9(8):1102–1123.

33. Hosein AN, Huang H, Wang Z, Parmar K, Du W, Huang J, Maitra A, Olson E, Verma U, Brekken RA. Cellular heterogeneity during mouse pancreatic ductal adenocarcinoma progression at single-cell resolution. JCI Insight 2019;4(16).

34. Bali MA, Metens T, Denolin V, Delhaye M, Demetter P, Closset J, Matos C. Tumoral and Nontumoral Pancreas: Correlation between Quantitative Dynamic Contrast-enhanced MR Imaging and Histopathologic Parameters. Radiology 2011;261(2):456–466.

35. Cantelmo AR, Pircher A, Kalucka J, Carmeliet P. Vessel pruning or healing: endothelial metabolism as a novel target? Expert opinion on therapeutic targets 2017;21(3):239–247.

36. White NS, McDonald C, Farid N, Kuperman J, Karow D, Schenker-Ahmed NM, Bartsch H, Rakow-Penner R, Holland D, Shabaik A, Bjornerud A, Hope T, Hattangadi-Gluth J, Liss M, Parsons JK, Chen CC, Raman S, Margolis D, Reiter RE, Marks L, Kesari S, Mundt AJ, Kane CJ, Carter BS, Bradley WG, Dale AM. Diffusion-weighted imaging in cancer: physical foundations and applications of restriction spectrum imaging. Cancer Res 2014;74(17):4638–4652.

37. Jensen JH, Helpern JA, Ramani A, Lu H, Kaczynski K. Diffusional kurtosis imaging: The quantification of non-gaussian water diffusion by means of magnetic resonance imaging. Magnetic Resonance in Medicine 2005;53(6):1432–1440.

38. Hempel JM, Schittenhelm J, Bisdas S, Brendle C, Bender B, Bier G, Skardelly M, Tabatabai G, Castaneda Vega S, Ernemann U, Klose U. In vivo assessment of tumor heterogeneity in WHO 2016 glioma grades using diffusion kurtosis imaging: Diagnostic performance and improvement of feasibility in routine clinical practice. Journal of neuroradiology = Journal de neuroradiologie 2018;45(1):32–40.

39. Zheng X, Carstens JL, Kim J, Scheible M, Kaye J, Sugimoto H, Wu CC, LeBleu VS, Kalluri R. Epithelial-to-mesenchymal transition is dispensable for metastasis but induces chemoresistance in pancreatic cancer. Nature 2015;527(7579):525–530.

40. Bhattacharjee S, Hamberger F, Ravichandra A, Miller M, Nair A, Affo S, Filliol A, Chin L, Savage TM, Yin D, Wirsik NM, Mehal A, Arpaia N, Seki E, Mack M, Zhu D, Sims PA, Kalluri R, Stanger BZ, Olive KP, Schmidt T, Wells RG, Mederacke I, Schwabe RF. Tumor restriction by type I collagen opposes tumor-promoting effects of cancer-associated fibroblasts. J Clin Invest 2021;131(11).

41. Shi M, Yu DH, Chen Y, Zhao CY, Zhang J, Liu QH, Ni CR, Zhu MH. Expression of fibroblast activation protein in human pancreatic adenocarcinoma and its clinicopathological significance. World journal of gastroenterology 2012;18(8):840–846.

42. Santos AM, Jung J, Aziz N, Kissil JL, Puré E. Targeting fibroblast activation protein inhibits tumor stromagenesis and growth in mice. The Journal of Clinical Investigation 2009;119(12):3613–3625.

